# Contribution of 3D genome topological domains to genetic risk of cancers

**DOI:** 10.1101/2021.07.26.453813

**Authors:** Kim Philipp Jablonski, Leopold Carron, Julien Mozziconacci, Thierry Forné, Marc-Thorsten Hütt, Annick Lesne

**Author notes:** Corresponding authors (MTH), (AL).

## Abstract

Genome-wide association studies have identified statistical associations between various diseases, including cancers, and a large number of single-nucleotide polymorphisms (SNPs). However, they provide no direct explanation of the mechanisms underlying the association. Based on the recent discovery that changes in 3-dimensional genome organization may have functional consequences on gene regulation favoring diseases, we investigated systematically the genome-wide distribution of disease-associated SNPs with respect to a specific feature of 3D genome organization: topologically-associating domains (TADs) and their borders.

For each of 449 diseases, we tested whether the associated SNPs are present in TAD borders more often than observed by chance, where chance (i.e. the null model in statistical terms) corresponds to the same number of pointwise loci drawn at random either in the entire genome, or in the entire set of disease-associated SNPs listed in the GWAS catalog. Our analysis shows that a fraction of diseases display such a preferential location of their risk loci. Moreover, cancers are relatively more frequent among these diseases, and this predominance is generally enhanced when considering only intergenic SNPs. The structure of SNP-based diseasome networks confirms that TAD border enrichment in risk loci differ between cancers and non-cancer diseases. Different TAD border enrichments are observed in embryonic stem cells and differentiated cells, which agrees with an evolution along embryogenesis of the 3D genome organization into topological domains.

Our results suggest that, for certain diseases, part of the genetic risk lies in a local genetic variation affecting the genome partitioning in topologically-insulated domains. Investigating this possible contribution to genetic risk is particularly relevant in cancers. This study thus opens a way of interpreting genome-wide association studies, by distinguishing two types of disease-associated SNPs: one with a direct effect on an individual gene, the other acting in interplay with 3D genome organization.

**Author summary:** Genome-wide association studies comparing patients and healthy subjects have evidenced correlations between diseases and the presence of pointwise genetic variations known as single-nucleotide polymorphisms (SNPs). We exploit and extend this statistical analysis by investigating the location of risk loci, i.e. disease-associated SNPs, with respect to the 3D organization of the genome into spatially-insulated domains, the topologically-associating domains (TADs).

We show that for certain diseases, mostly cancers, their associated risk loci are preferentially located in the borders of these topological domains. The predominance of cancers among these diseases is confirmed and even enhanced when considering only intergenic SNPs. A different enrichment behavior is observed in embryonic stem cells and derived cell lines at an early developmental stage, presumably due to the not fully mature TAD structure in these cells.

Overall, our results show that genome variations in specific TAD borders may increase the risk of developing certain diseases, especially cancers. Our work underlines the importance of considering the genetic risk loci within their 3D genomic context, and suggests a role of 3D genome partitioning into topological domains in the genetic risk which differs between cancers and non-cancer diseases.

## INTRODUCTION

Genome-wide association studies (GWAS) have compared the genomes of large cohorts of patients and healthy individuals, and evidenced statistical associations between the presence of single-nucleotide polymorphisms (SNPs) in their variant form, and the presence of diseases [1,2], including cancers [3,4]. Remarkably, only a few percent of disease-associated SNPs (daSNPs) are located in coding regions of the genome [5,6]. How the vast majority of non-coding daSNPs are mechanistically related to the risk of developing a disease is yet unclear. While SNPs were at first considered as mere markers of the nearest gene, it rapidly appeared that they can have a direct functional role in affecting the regulation of neighboring genes, typically by being located in regulatory sequences [6,7] or non-coding RNA sequences [8]. Systematic analyses of SNP location with respect to genome annotations such as binding sites of regulatory proteins or histone epigenetic marks correlated with promoter or enhancer loci [9-11], as well as joint transcriptome analysis [12], have been successfully used to identify causal variants and affected genes.

However, such correlation analyses are not sufficient to unravel all the determinants of SNP-disease associations. An additional ingredient, the 3-dimensional (3D) genome organization, must be taken into account. Indeed, it is now acknowledged that not only the adjacent sequences of the gene, but also the 3D folding of the genome play a role in genomic functions [13], and more specifically in the regulation of gene expression [14], its evolution [15], and its misregulation e.g. in cancers [16]. It thus appears essential to reformulate the notion of genetic risk to a disease in the context of the recently acquired knowledge about 3D genome organization. Our working hypothesis is that certain non-coding SNPs could, in their variant form, affect the 3D genome architecture and its role in gene regulation, thus favoring the development of diseases.

This idea has been documented experimentally for enhancer-promoter loops [17,18]. A variant at a single SNP may induce a change in the looping bringing enhancer and promoter into close spatial proximity, henceforth affecting the expression of the corresponding genes. For instance, a single SNP may modify a CTCF binding site and in turn nucleosome positioning and chromatin 3D architecture, a documented situation for asthma risk [19,20]. At the MYB locus, 3C experiments have shown reduced interactions between promoter and enhancer in the presence of the at-risk allele, providing an instance of a SNP having a causal architectural effect [21].

Pursuing this line of investigation at the genome-wide level, we will consider another feature of 3D genome structure, namely topologically-associating domains, TADs [22]. There are few genomic contacts between two adjacent TADs, and the insulating capacity of TAD borders [23] has been shown to be essential for proper gene regulation, by preventing spurious interactions between genes and enhancers located in adjacent TADs [24-26]. Gene misregulation can occur due to TAD border disruption induced by the presence of short tandem repeats [27]. The importance of TADs and the effect of TAD border disruption (dashed arrow in Fig 1) has also been shown in the case of *Hox* genes [28, 29], or as a way to control developmental genes in drosophila embryos, as we recently proposed [30]. An effect of TAD border disruption has also been observed in cancer cells as a consequence of cancerous mutations [31,32].

**Fig 1.**
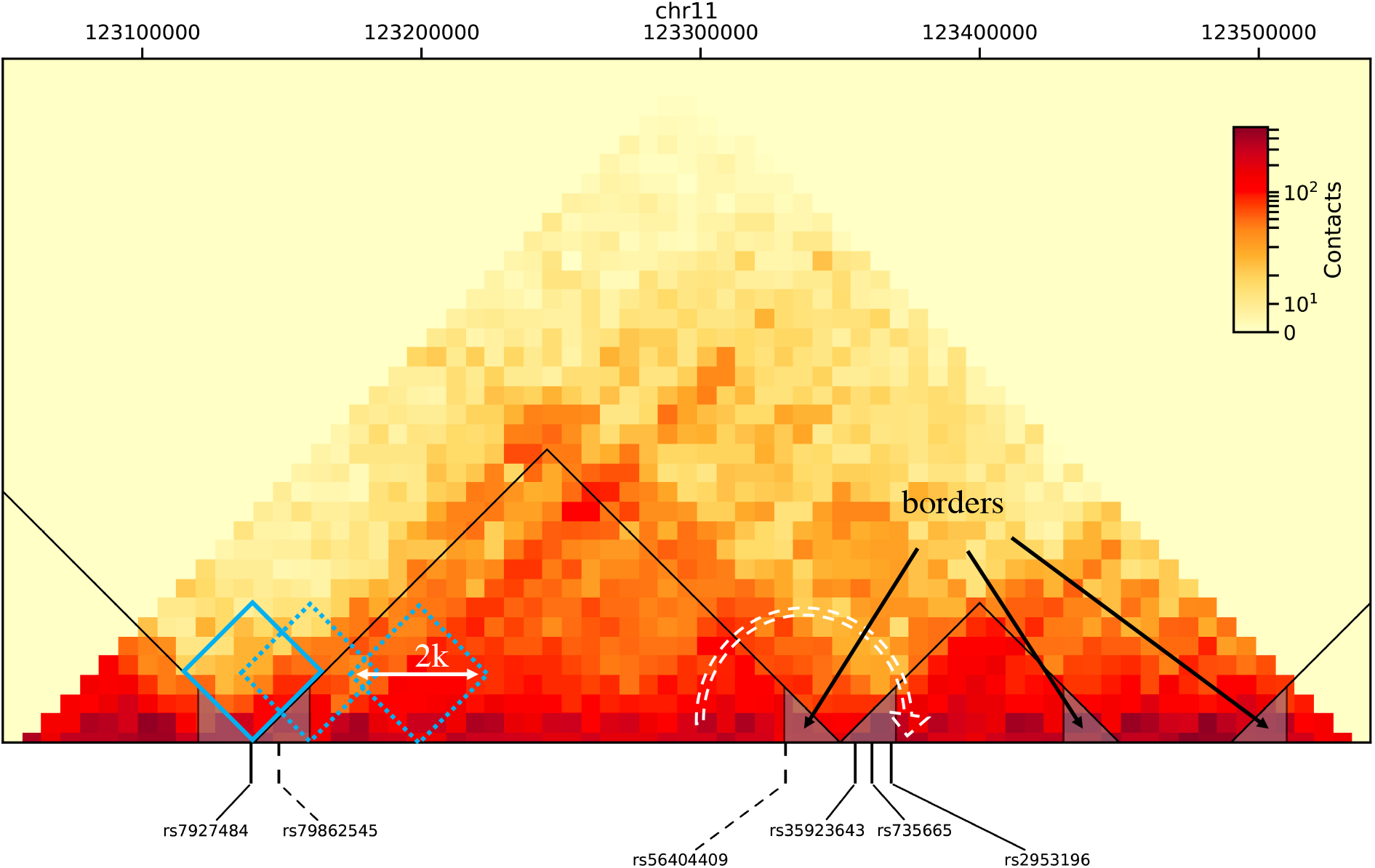
Schematic of the investigation. Underlying Hi-C data are displayed as a heat map (the redder the more contacts, as indicated in the color bar), here for a region of chromosome 11 (chr11: 123050000-123550000, hg19 coordinates), drawn from data published in [34], for IMR90 cell type, at 10kb-resolution. TADs are underlined with black triangle lines. They are determined using TopDom algorithm [35], which identifies a demarcation between two TADs as a local minimum of the number of contacts in a sliding window of size 2*k* bins, where *k* is a tunable parameter (blue diamonds, the full-lined one corresponding to the limit of a TAD). TAD borders are defined as 20kb-regions from TAD ends inward, and underlined here as small triangles filled in grey. Vertical lines pinpoint disease-associated SNPs located in TAD borders (full line for cancer-associated SNPs, dashed lines for SNPs associated with non-cancer diseases). For each cancer or each non-cancer disease, we investigate the potential over-representation of its associated SNPs in TAD borders. The dashed white arrow indicates increased physical contacts and regulatory interactions between adjacent TADs that would possibly appear in a border affected by the presence of an at-risk SNP allele.

The guideline of our computational study is that functional mechanisms underlying the association of risk loci with a disease may, in some cases, be mediated by a change occurring in TAD borders when embedded SNPs are in their at risk-form. Accordingly, the increased disease risk due to the presence of non-coding SNPs in their variant form could presumably be better understood by investigating their location with respect to various features of the 3D genome organization, in particular its partitioning into TADs [26].

We studied quantitatively and systematically, for each disease, the location of disease-associated SNPs with respect to TAD borders (Fig 1). Different cell types and data obtained in different laboratories have been considered (overall 15 datasets). Based on a preliminary analysis [33] and on their different etiology, we analyzed separately 71 cancers and 378 non-cancer diseases and compared the distribution of their associated SNPs with respect to TAD borders. We analyzed, for each disease, the distribution of its disease-associated SNPs in TAD borders compared to chance, where chance corresponds to the same number of pointwise loci drawn at random either in the entire genome or in the set of SNPs listed in the GWAS catalog. We investigated whether the results persist when considering only intergenic SNPs (40% of the total set of daSNPs, overall comprising 21,183 entries). An integrated pipeline, sketched in S1 Fig, has been devised for this analysis, and its different elements are detailed in the *Materials and methods* section.

We emphasize that the present investigation does not consider somatic mutations appearing in cancer cells along cancer progression. It focuses on genomic variations observed from birth in the genome of any healthy cell of the individual.

## MATERIALS AND METHODS

### Disease-associated SNPs (daSNPs)

We used the version v1.0.2-associations_e94_r2018-09-30 of the *GWAS catalog*: www.ebi.ac.uk/gwas/ [36,37]. We extracted all SNP entries associated with a disease EFO term (*Experimental Factor Ontology*), overall 449 EFO terms, distinguishing 71 cancers and 378 non-cancer diseases. Specifically, diseases were found by selecting all traits which fall into the disease subtree (EFO_0000408) from the EFO ontology, and subsequently cancers are separated from non-cancer diseases by using the cancer subtree (EFO_0000311).

A SNP can be associated with a disease multiple times in the GWAS catalog when it was found in distinct studies; we ignored this multiplicity by dropping duplicates of its identifier *snpId*.

We classified the resulting 21,183 daSNPs into intergenic (40%), intronic (55%) or exonic (5%) type according to its parent category in *The Sequence Ontology* database, (http://www.sequenceontology.org), version 2015-11-24, [38].We then identified for each disease the subset of its associated intergenic SNPs (on average 47 daSNPs and 18 intergenic daSNPs for both cancer and non-cancer diseases).

### Hi-C data

We used the *Cooler* Hi-C database [39] at ftp://cooler.csail.mit.edu/coolers, which provides published Hi-C data files in the .cool format, at 10kb-resolution (bin size). Throughout our study, genomic coordinates refer to the hg19 genome version adopted in this database. We present in the main text results obtained with high-resolution data from E. Lieberman-Aiden’s laboratory [34] for the 5 native and non-cancerous cell lines available, namely GM12878 (human lymphoblastoid cell line, data obtained with MboI or DpnII restriction enzyme), IMR90 (fetal lung fibroblasts of Caucasian origin), HMEC (human mammary epithelial cells), NHEK (normal human epidermal keratinocytes) and HUVEC (human umbilical vein endothelial cells). We did not consider cancer cell types as we are interested in the genetic risk present at birth in all cells. We also investigated datasets from B. Ren’s laboratory [22, 40-42], obtained in pioneering experiments using a lower sequencing depth and an enzyme HindIII producing larger restriction fragments (see S2 Fig), for several cell types: GM12878 in [41], IMR90 in [22,40] embryonic stem cells (H1 hESC) in [22,42], and cell lines derived from H1 hESC in [42], namely mesendoderm (H1_ME), neural progenitors (H1_NP) trophoblast-like cells (H1_TB) and mesenchymal cells (H1_MS). Overall 15 datasets were examined in our study.

### TAD determination

We determined TAD coordinates using the *TopDom* algorithm [35], applied after a transformation of .cool files into count matrices using https://github.com/open2c/cooler. Its principle is to count the number of contacts in a window sliding along the genome and to locate TAD ends at the minima of this count (see Fig 1, blue diamonds). Genomic regions having established only very few contacts in the experiment, labelled ‘gap’ by TopDom, were filtered out. The choice of using this algorithm is supported by comparative studies [43-45]. Moreover, TopDom is based on quantifying the topological insulation between adjacent TADs, which is the feature that matters for gene (mis)regulation. In particular, recently evidenced long-range associations between TADs and their higher-order organization [46] will not be considered here.

The TAD caller thus involves a tunable parameter *k*, measuring the half-size (in bins, of length equal to the chosen resolution) of the sliding window. This parameter offers a way to investigate the well-known variability in TAD determination [43-45], as depicted in S3 Fig. The general trend is that larger numbers of TADs are observed for lower values of *k*, at which substructures are also extracted while only large TADs are extracted by the algorithm at large values of *k*. To overcome this technical variability, we scanned all values of the window size *k* from *k*=3 to *k*=20 and adopted two strategies: either (i) to aggregate our observations over these values of *k* (Fig 2), or (ii) to use a more stringent majority rule in further analyses (Figs 3, 4). Both strategies reduce small-number effects and smooth out TAD variability, overall yielding robust results despite the lack of robustness of the TAD landscape (S3 Fig and S4 Fig).

**Fig 2.**
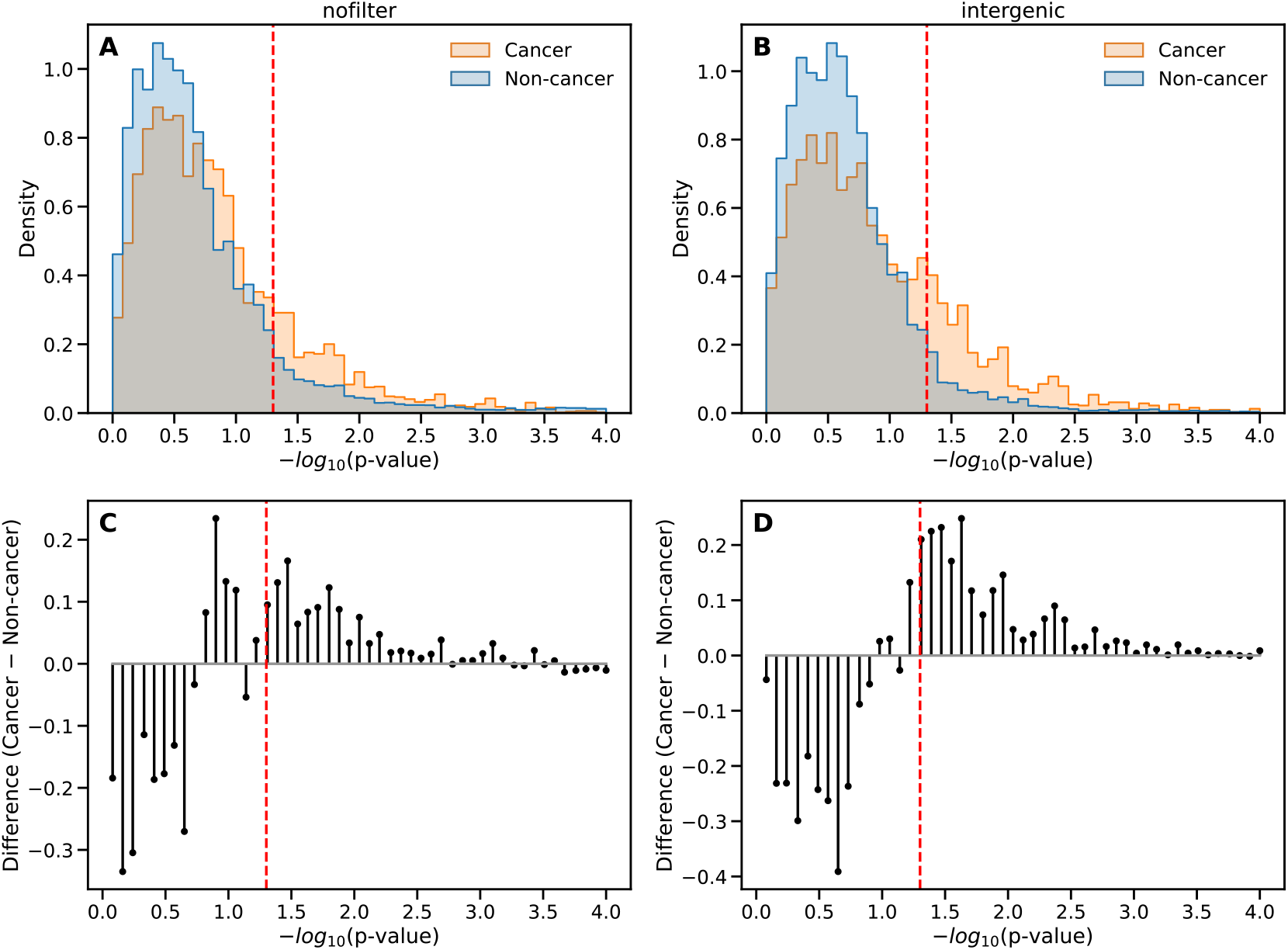
Preferential location in TAD borders of the SNPs associated to a disease. **(A)**Normalized histogram of enrichment *p*-values (corrected for multiple testing and plotted as -log_10_ *p* on the horizontal axis) for cancers (yellow bars) and non-cancer diseases (blue bars, overlap in grey), considering the contribution of all SNPs to a potential TAD border enrichment. The results have been aggregated over datasets for different cell types from [34] at 10kb-resolution and all considered values of the window parameter *k* of TopDom algorithm. The two histograms have been normalized separately. The dashed red line indicates the significance threshold at *p\**=0.05. **(B)** Same as (A) considering only intergenic daSNPs in the hypergeometric enrichment test. (**C**) Difference between the cancer histogram, in orange, and the histogram for non-cancer diseases in (A), showing that relatively more cancers display a preferential location of their associated SNPs in TAD borders. (**D**) Same as (C) for the histograms in (B), showing that the relative dominance of cancers, among diseases displaying TAD border enrichment, is enhanced when considering only intergenic daSNPs.

**Fig 3.**
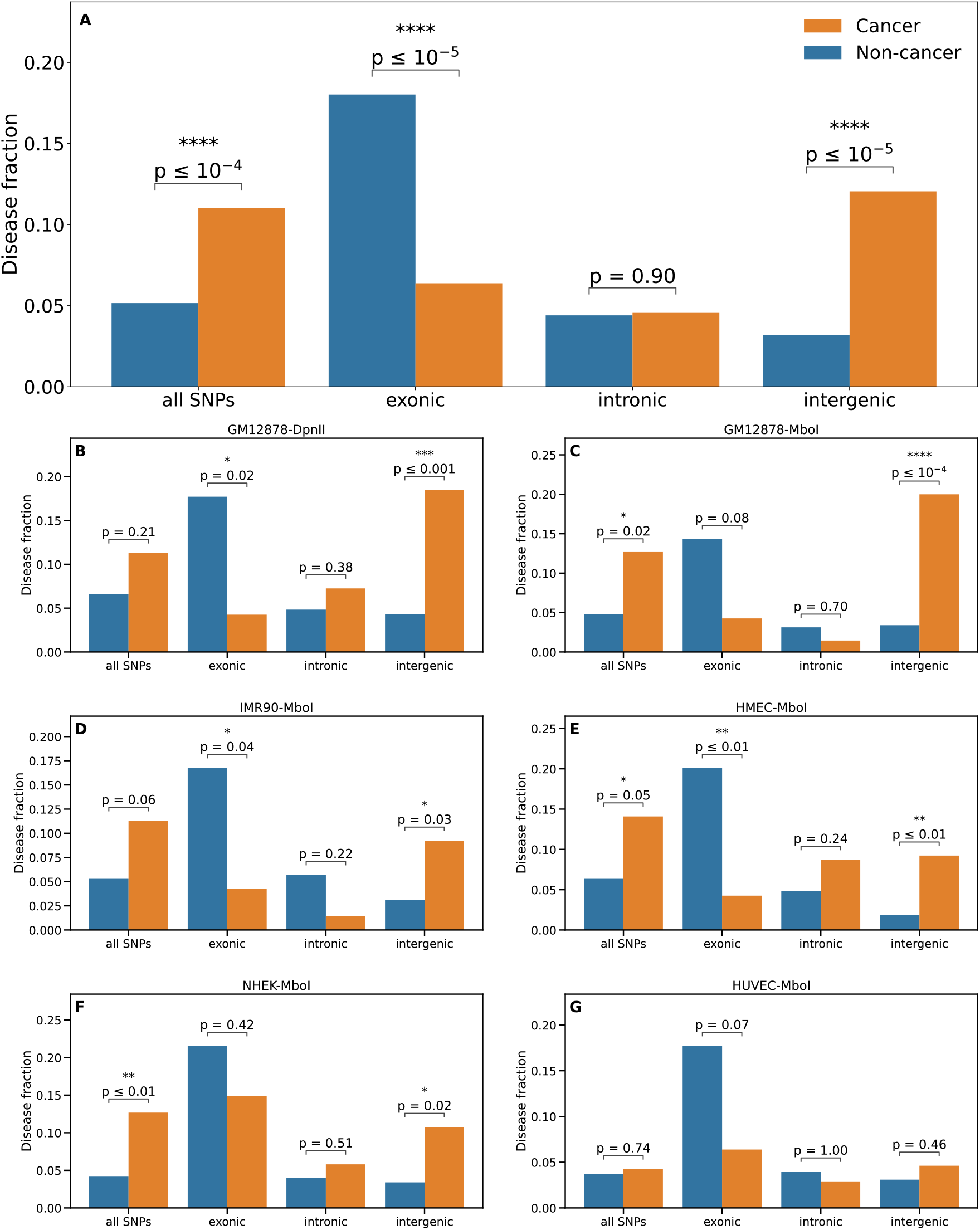
Preferential location of daSNPs in TAD borders occurs relatively more often for cancers, and this feature is generally enhanced for intergenic daSNPs. A comparison of the fraction of cancers (orange) and fraction of non-cancer diseases (blue) displaying TAD-border enrichment is presented for various filters on the GWAS catalog: considering all daSNPs, then only exonic, intronic, or intergenic daSNPs. Only diseases displaying enrichment for a majority of values of the TopDom window-size parameter *k* are included (see *Materials and methods*). **(A**) Aggregation over 5 cell types (6 datasets) using data at 10kb-resolution from [34]. (**B-G**) Detailed comparison for each of the 6 datasets, where the cell type is indicated above each panel, together with the restriction enzyme (DpnII or MboI) used in the Hi-C experiment. Stars indicate when the difference between cancers and non-cancer diseases is statistically significant (Fisher exact test, *: *p*≤0.05, **: *p*≤0.01, ****: *p*≤0.0001). Analyses for data from [22, 40-42] are presented in S8 Fig.

**Fig 4.**
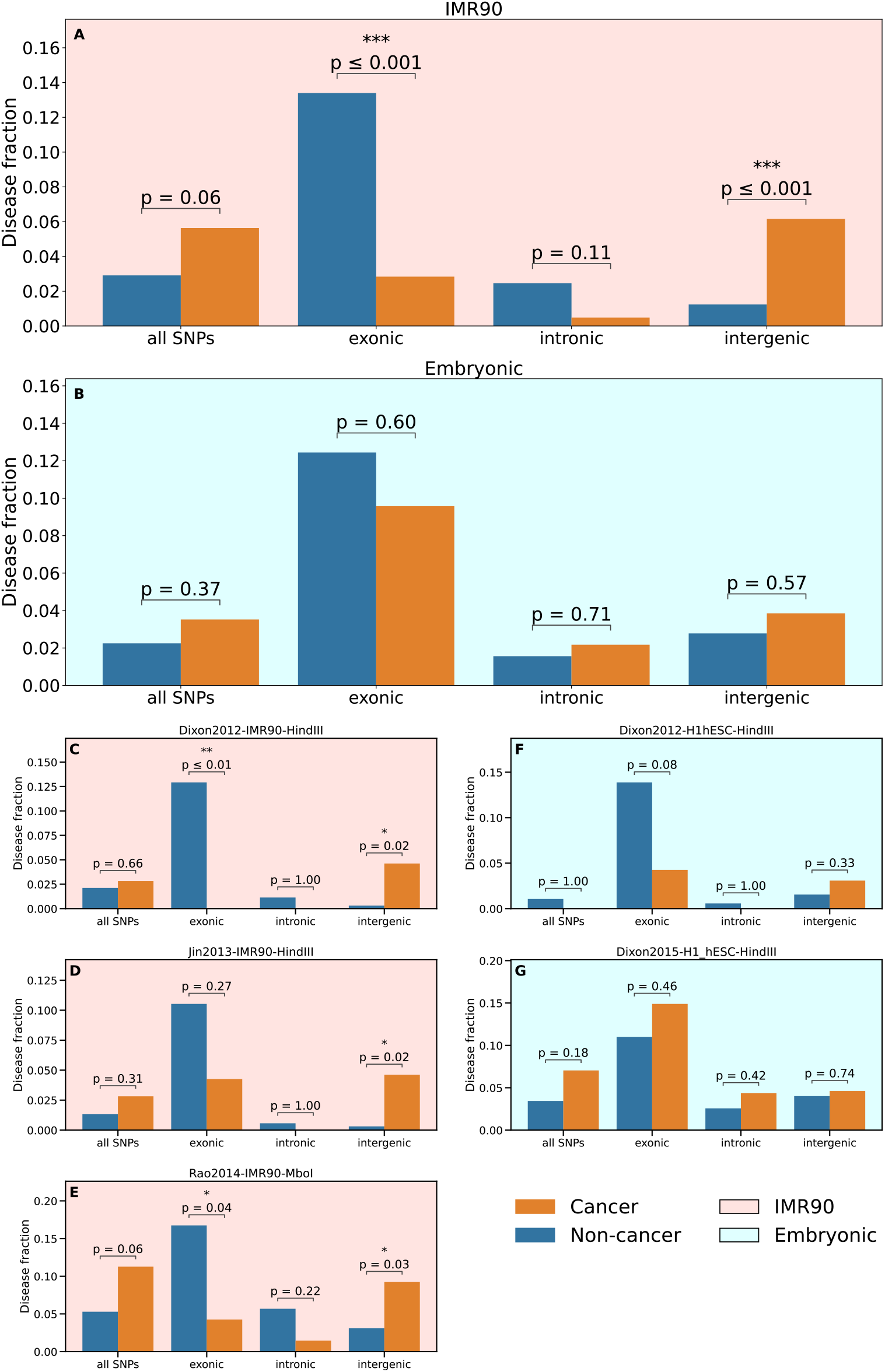
Comparison of enrichment results for IMR90 cells and embryonic cells. Same as Fig. 3, now comparing three datasets for IMR90 cell line (left column) with two datasets for embryonic stem cells (hESC). The two panels at the top display the results aggregated over the individual datasets, for IMR90 cells (pink background) and stem cells (blue background), respectively.

A discrepancy between TADs determined with TopDom and the visual impression given by the contact map could appear locally (see e.g. Fig 1), coming from the following difference: TopDom is based on the insulation of TADs, i.e. the presence of low-density (yellow) zones between the triangles delineating the TADs (see Fig 1), whereas the alternative understanding of a TAD as a region with an increased density of internal contacts would rather focus on dark red triangles emerging from the background in the Hi-C map.

### TAD borders

We defined *TAD borders* as regions of 20kb located *inside* the TAD at the limits of this TAD. Their size has been chosen smaller than the median size (23kb) of a human gene [47].

This definition agrees with a topological characterization of TAD borders as regions across which the contact frequency displays a marked decrease [48] and is consistent with the use of TopDom for calling TADs. It differs from the notion of *TAD boundary*, considered e.g. in [22,27] which is the ---not always existing--- linker region *between* two successive TADs along the genome (not belonging to any TAD), whereas a TAD will always have two borders. TAD borders cover from 8% up to 14% of the genome when TopDom parameter *k* varies, the smallest fraction being observed for *k*=20 (see S1 File, S1 Table and S2 Table in *Supporting Information*)

### Preferential location of daSNPs in TAD borders (TAD border enrichment)

For each disease (EFO term), we have tested whether the associated SNPs are located in TAD borders more often than observed by chance, where chance (i.e. the null model in statistical terms) corresponds to the same number of pointwise loci drawn at random in the entire genome. In a second analysis, chance corresponds to the same number of SNPs drawn at random in the entire set of disease-associated SNPs listed in the GWAS catalog. The statistical significance of a preferential location of daSNPs in TAD borders is then assessed by computing a *p*-value for the disease according to a hypergeometric test.

In more detail:

Testing a preferential location in TAD borders of the SNPs associated to a given disease involves the hypergeometric distribution *H (q* | *N, n, Q)* where *Q* is the number of SNPs associated to the considered disease and *q* the part located in TAD borders. In the *genome-based null model, N* is the total number of base pairs in the genome and *n* the part located in TAD borders. In practice, given the finite resolution which amounts to discretize the genome in 10kb-bins, *N* is the total number of bins in the genome and *n* the number located in TAD borders. In the *SNP-based null model*, close to that considered e.g. in [49], *N* is the total number of disease-associated SNPs listed in the GWAS catalog (each counted once, and not including the SNPs associated with non-pathological traits so that the random model is the closest possible to the data) and *n* the part located in TAD borders. Results presented in the main text were obtained with the more conservative genome-based null model. A comparison with those obtained with the SNP-based null model is presented in S5 Fig.

The *p*-value for the considered disease, assessing the over-representation of its associated SNPs in TAD borders, is computed as the cumulative distribution function (i.e., the fraction of values larger than or equal to *q*) of this hypergeometric distribution. Given the symmetry *H (q* | *N, n, Q) = H (q* | *N, Q, n*) of the hypergeometric distribution, it is equivalent to state that (i) the SNPs associated to the disease are located in a TAD border more often than expected by chance, or (ii) TAD borders contain a SNP associated to this disease more often than expected by chance. We term such a situation *TAD border enrichment*.

After computing a raw *p-*value for each disease, Benjamini-Hochberg procedure (*multipletests* function with method *fdr_bh* from the *statsmodels* package in Python, www.statsmodels.org/dev/index.html#citation) is applied to obtain *p*-values adjusted for multiple testing, so that the false discovery rate is controlled at level 5% when the adjusted *p*-value is lower than 0.05 [50]. Given the overwhelming number of non-cancer diseases (378) compared to cancers (71) and their different etiology, we investigated separately these two groups of diseases. These two groups are well-defined on biological criteria independently of our enrichment testing, so that correction for multiple testing has been applied separately in each group. We checked that our main result (the relatively more frequent TAD border enrichment for cancers) is also qualitatively observed when using a global multiple-testing correction, or even no correction (S6 Fig and S7 Fig).

### Enrichment histograms

Histograms of corrected *p-*values (Fig 2) are plotted and normalized separately for cancers and non-cancer diseases. The counts have been first aggregated over the 6 considered datasets from [34] and the values of the window parameter *k* of the TAD caller. In order to get a better display of the core features of the plots, the range of *p*-values has been truncated at log_10_(1/*p*) = 4. A few EFOs, with corrected *p-*values smaller than 0.0001, are thus lying outside the displayed plot. It would be possible to choose a larger bin size or even to draw a smooth histogram, but this would rather dilute the information.

### Comparison of TAD-border enrichment for cancers and non-cancer diseases

Due to the small number of cancers (71) compared to non-cancer diseases (378), we compared the fraction of cancers and the fraction of non-cancer diseases displaying a significant TAD border enrichment, considering either all their associated SNPs or only a sub-category (exonic, intronic and intergenic). Only diseases displaying a significant enrichment for a majority (more than 50%) of values of the window parameter *k* have been counted. The significance of a difference between the disease fractions has been assessed using Fisher’s exact test. The comparison has been done for each of the 6 considered datasets (for 5 cell types) from [34], and also after aggregating the disease counts over these datasets.

### Workflow

The different steps described above have been gathered in an easy-to-execute pipeline, using *Snakemake*

(https://snakemake.github.io, [51]) unifying the analysis of different datasets. Its rule graph is presented on S1 Fig.

Its code, written in Python, is freely available at https://github.com/kpj/GeneticRiskAndTADs. An order of magnitude of the typical numbers of diseases and SNPs of different categories involved in our analysis is provided in *Supporting Information* (S1 File, S1 Table, S2 Table).

### SNP-based diseasome network and its analysis

We introduced a network representation where nodes are diseases and an edge is drawn between two diseases when they share an associated SNP. Starting from a bipartite network relating diseases and their associated SNPs, this representation is the projection on disease nodes. It is the analogue for SNPs of the network relating diseases and their associated genes, known as the diseasome, and its projected version [52]. A filter has been applied on shared SNPs: in the network labeled ‘border SNPs’, an edge is drawn when the diseases share a SNP lying in a TAD border for a majority of values of the window parameter *k* (underlying Hi-C data from [34], IMR90 cell type). The network labeled ‘non-border SNPs’ involves the complementary set of SNPs. Additionally, the two networks have been re-drawn considering only intergenic SNPs. Non-cancer disease and cancer nodes (and among them, those displaying TAD border enrichment) were underlined with different colors. Note that each cancer or each non-cancer disease is associated with an ensemble of border SNPs, of non-border SNPs, of intergenic border SNPs and of intergenic non-border SNPs, and could be present in more than one of the four networks. In Fig. 5, the four networks have been visualized using *NetworkX* Python package. For each network, only diseases (nodes) having an associated SNP of the prescribed type are drawn.

**Fig 5.**
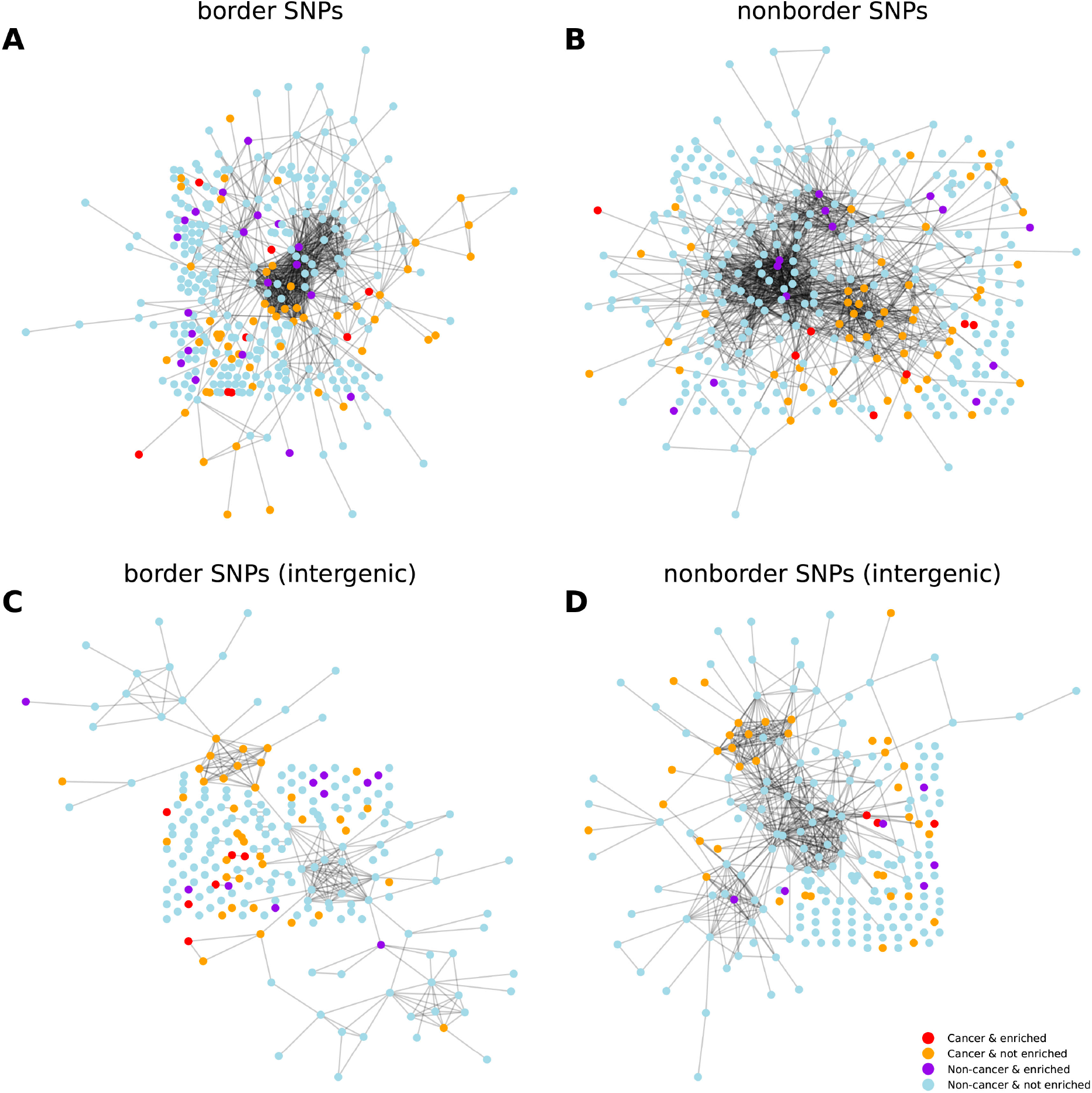
Clustering of cancers and non-cancer diseases sharing daSNPs. In this network representation, red (resp. violet) nodes correspond to cancers (resp. non-cancer diseases) displaying TAD border enrichment in daSNPs, orange (resp. light blue) nodes to other cancers (resp. other non-cancer diseases). An edge is drawn between two diseases when they share an associated SNP (A) located in a TAD border for a majority of values of the window parameter *k (*border SNPs, see *Materials and methods)* or (B) belonging to the complementary set (non-border SNPs). (C, D) Same as (A, B) when considering only intergenic SNPs. The four networks have been visualized using *NetworkX* Python package. Underlying Hi-C data from [34], IMR90 cell type, 10kb resolution.

For a quantitative comparison of the four networks regarding their clustering and assortativity properties, we computed an indicator for any subset of nodes, e.g. a group of nodes with the same color. This indicator, called *network coherence*, is defined as the z-score of the number of edges within the subset of nodes, compared to a thousand randomly drawn groups of nodes of the same size [53,54]. Network coherence thus measures whether the induced subgraph is more densely connected (i.e. contains more links) than expected at random in the original network. As a z-score, it provides an absolute quantification, independent of the overall size of the group, which makes cross-comparisons possible. Choosing a threshold larger than 1 on the number of shared SNPs required to draw an edge does not change qualitatively the results but reduces the number of diseases involved, which brings statistics to a limit. All the results presented in the text were obtained with a threshold equal to 1.

## RESULTS

### 1. For a fraction of diseases, their associated SNPs are preferentially located in TAD borders

We used 15 high-resolution published Hi-C datasets, obtained in two different laboratories, for different human cell types and using different restriction enzymes. We first determined the 3D genome organization in topological domains (TADs) using TopDom algorithm. Then for each of the 449 diseases considered, the potential over-representation of its associated SNPs in TAD borders has been assessed using a hypergeometric test (see *Materials and methods*).

Results are shown in Fig 2A in the form of a *p*-value histogram. The *p-*value for a given disease measures the statistical significance of the preferential location of its associated SNPs in TAD borders, or equivalently, the statistical significance of TAD-border enrichment in its associated SNPs. Given the overwhelming number (378) of non-cancer diseases compared to cancers (71) present in the GWAS catalog, and their different etiology, we considered separately cancers and non-cancer diseases.

Our analysis shows that for a fraction of diseases, their associated SNPs are preferentially located in TAD borders, as already observed in a preliminary study [33]. A recent study centered on complex trait heritability consistently put forward a specific status of TAD borders [55]. In our results, the fact that only a fraction of diseases display such a TAD border enrichment precludes a trivial explanation related to some confounding feature of the TAD borders, that would produce enrichment for all diseases.

Despite the well-known variability in TAD determination [43-45] (S3 Fig, S4 Fig), we get a clear statistical result, both for a single value of the parameter *k* (window size) of the TAD caller TopDom, or when aggregating over several values of this parameter.

### 2. TAD-border enrichment in daSNPs is yet observed with intergenic SNPs alone

To better interpret the over-representation of daSNPs in TAD borders, we considered specific subsets of daSNPs according to their genomic location: namely daSNPs located in exons (5%), whose variant form has an impact on a protein sequence (coding SNPs) and those, by far more common, located either in introns (55%) or in intergenic regions (40%). We then performed the enrichment analysis considering only intergenic daSNPs. Fig 2B shows that TAD border enrichment is still observed and even enhanced for intergenic SNPs.

### 3. Preferential location of daSNPs in TAD borders is observed mostly for cancers

The normalized histograms presented in Fig 2A indicate that the fraction of cancers displaying TAD border enrichment in daSNPs is larger than the fraction of non-cancer diseases displaying such a preferential location of their associated SNPs. To better evidence this relative predominance of cancers, we present on Fig 2C the difference between the two histograms. Considering a percentage among cancers is motivated by the small number of cancers (71) in the whole set of diseases (449 EFOs listed in the GWAS). An alternative and equivalent formulation is that the fraction of cancers among the diseases displaying enrichment is larger than the fraction of cancers among the overall set of diseases.

This observation motivated a systematic quantification of the relative predominance of cancers, presented in Fig 3 together with an assessment of its statistical significance. Similar results have been obtained for the genome-based and the SNP-based null models (see *Materials and methods*), as expected from the comparison of the enrichment *p*-values obtained with these two null models, presented in S5 Fig.

We analyzed further the robustness of this relative predominance of cancers among the diseases displaying TAD border enrichment. We observed that it is sensitive but overall robust with respect to the parameter *k* of the TAD caller (S3 Fig). This relative predominance of cancers is equally observed before correcting for multiple testing or using a global correction (see S6 Fig), demonstrating that it is not an artefact of applying the correction separately for cancers and non-cancer diseases.

### 4. The relative predominance of cancers among diseases displaying TAD-border enrichment is enhanced when considering intergenic SNPs alone

We then dissected the contribution of exonic, intronic and intergenic SNPs to TAD border enrichment. Mainly, the relative predominance of cancers among diseases displaying TAD border enrichment in intergenic daSNP is confirmed and even enhanced compared to the quantification involving all types of daSNPs (Fig 2D and Fig 3A). The same shift is observed for both null models (data not shown, but see S5 Fig), and it is robust with respect to the type of multiple-correction adopted (see S7 Fig).

That exonic SNPs also contribute to TAD border enrichment in daSNPs is not surprising nor contradictory, as TAD borders are known to host active genes [22,56]. We actually observe a stronger and unexpected feature, namely a relative dominance of non-cancer diseases when restricting to exonic daSNPs. However, the low number (a few units) of exonic daSNPs for most diseases brings statistics to a limit, and precludes elaborating too much on this observation. In particular, although visible in all cases, the relative dominance of non-cancer diseases reaches statistical significance only in the aggregated data. This observation nevertheless confirms the specificity of cancers, compared to non-cancer diseases, regarding the role of TADs in the associated genetic susceptibility.

### 5. Different enrichments are observed in embryonic stem cells (hESC)

We consistently observe the above-described results for most cell types, as shown in Fig 3B-G and S8 Fig. The relative dominance of cancers among diseases displaying TAD border enrichment in their associated intergenic SNPs largely reaches statistical significance, with a few notable exceptions: umbilical vein cells (HUVEC, Fig 3G), embryonic stem cells (hESC, S8 Fig A, E). H1-derived cells (H1_ME, H1_NP, H1_TB, S8 Fig F, G, H).

We further assess the different enrichment results between embryonic stem cells and differentiated cell types in a systematic comparison of hESC and IMR90 cell lines, for which several datasets from different studies are available, Fig 4. Consistent results are obtained for the two datasets obtained in hESC (Fig 4, blue panels), and the three datasets for IMR90 (Fig 4, pink panels), respectively. These results can be extracted more clearly by an aggregation over datasets for the same cell type (Fig 4, top panels). The comparison emphasizes the peculiarity of hESC, with no relative predominance of cancers among diseases displaying TAD border enrichment in intergenic daSNPs. In H1 hESC and derived cells at early developmental stage (H1_ME, H1_NP, H1_TB), 3D genome partitioning in TADs is not significantly related to the location of cancer risk loci. Consistently, H1_MS cell line differs from the other H1-derived cell lines both in our analysis and in the original study [42], which has evidenced a genome-wide evolution of TAD structure in this series of H1-derived cell lines following their progressive differentiation. We also observe that HUVEC line has the same signature as hESC, with no cancer-specific features in the location of at-risk SNPs with respect to the TAD structure present in this cell line.

Our analysis thus suggests that TAD border contribution to the genetic risk of cancers is not observable in HUVEC, hESC and derived cells at early developmental stage (H1_ME, H1_NP, H1_TB) possibly due to their different (not fully mature) TAD structure [42, 57].

### 6. SNP-based diseasome analysis shows the unimportance of daSNPs sharing in the observed features

Among the disease-associated SNPs listed in the GWAS catalog, about 14% are actually associated to several diseases. To analyze the influence of such events on our results, we devise a network representation, where the nodes of the network are diseases (coloring differently cancers and non-cancer diseases) and a link is drawn between two diseases when they share at least one associated SNP. This is nothing but the SNP-based analog of the diseasome networks introduced in [52], in which a link is drawn between two diseases when they share a related gene. We then compared the networks obtained when a link is drawn only when the diseases share a border SNP, i.e. a SNP located in a TAD border for a majority of values of the window parameter (Fig 5A), when the diseases share a SNP belonging to the complementary set (non-border SNPs, Fig 5B), then with an additional filter keeping only intergenic SNPs (Fig 5C-D).

We observe that cancers share SNPs preferentially with other cancers, while non-cancer diseases share daSNPs preferentially with other noncancer diseases, corresponding in the network language to an assortative clustering of cancer nodes and non-cancer nodes, respectively. Similar clustering properties are observed in the networks based on border SNPs or intergenic border SNPs.

As the visual inspection of the networks can be misleading, we quantified for various node subsets the statistical enrichment of links of each induced subgraph. For this purpose, we used an indicator termed the *network coherence* [53,54]. Network coherence basically assesses whether the nodes in the set under consideration are more (or less) connected to each other than expected at random. It is defined as a z-score (see *Materials and Methods*) so as to get absolute values than can be used for comparing networks. A random subgraph would have a vanishing network coherence, while a positive value indicates a significantly higher number of internal links compared to random sets of the same size. S3 Table lists all network coherences for all four networks and all six node types: cancer or non-cancer, displaying TAD border enrichment (termed in short ‘enriched’) or not.

For non-cancer diseases the enrichment status makes a difference: The passage from not-enriched to enriched leads to a change of sign (from negative to positive) in the network coherence, suggesting a more important contribution of specific shared SNPs to TAD border enrichment in the class of non-cancer diseases.

For cancers the enrichment status does not make a difference: Both subsets of cancers (not enriched and enriched) have a high network coherence and hence, qualitatively speaking, display similar overlaps among SNP sets, either border SNPs or non-border SNPs. The presence of SNPs associated with multiple cancers therefore has a similar impact on our analysis both for cancers displaying TAD border enrichment and for other cancers. Moreover, the effect becomes weaker when imposing the additional filter of intergenic SNPs. Therefore, our statistical observations regarding TAD border enrichment in cancer-associated SNPs do not arise from a few shared border SNPs but actually from their preferential location in TAD borders.

## DISCUSSION

We have explored the notion of genetic risk to a disease in the context of the recently acquired knowledge about 3D genome organization, specifically the genome partitioning into topological domains (TADs). We have provided statistical evidence that for some diseases, mostly cancers, associated SNPs are preferentially located in TAD borders. Cancers are relatively more frequent among these diseases, and this relative predominance is enhanced when considering only intergenic SNPs. Network analysis demonstrates that these results are not due to a small number of SNPs common to several diseases.

The fact that the associated SNP is not necessarily the causal variant and may be only a marker related to the causal variation through linkage disequilibrium, does not affect significantly our conclusion: the correlated variations would still be located in the corresponding TAD border, whose size (20kb) is typically larger than the range of linkage disequilibrum. A wider genetic variation (more extended along the genome than a single point mutation) would even make more plausible an involvement of architectural mechanism in the risk of developing the disease.

While TAD organization in cancer cells is globally largely intact, many studies have shown that chromatin architecture can be disrupted in cancer by changes in TAD boundaries due to the vast genetic alterations, including copy number variation, mutations, translocations that accompany cancer development and progression (reviewed e.g. in [32]). Such disruptions lead to aberrant gene expression within the affected TADs. However, very little is known about how genetic SNP variants associated with cancer susceptibility are favoring cancer development in healthy individuals (i.e. before cancer development) and how they impact 3D genome organization. Using capture Hi-C approaches, it was shown that GWAS SNP variants associated with cancer in breast [58], prostate [59] and in colorectal tissues [60] affect long-range chromatin interactions. These pioneer studies suggested that such genetic variants drive altered expression of certain oncogenes and tumor suppression genes, but their impact was so far restricted to chromatin loop organization within TADs. Our results suggest a different mechanism of genetic risk for cancers and non-cancer diseases. Specifically, cancers might be promoted by joint mis-regulation of oncogenes through a weakening of TAD borders, while non-cancer diseases would rather be favored by local mis-regulations of specific genes.

The weaker relationship between genetic risk loci and TAD borders observed in hESC is consistent with the experimental evidence in mammals of a progressive maturation of the internal structure of TADs, with the establishment of additional enhancer-promoter interactions and further sub-TAD structures during cell differentiation [42, 57]. In other cell lines investigated, TAD border enrichment in intergenic daSNPs discriminates cancers and non-cancer diseases and suggests an essential difference about the role of genome organization in TADs as regards their genetic risk. These results will now have to be confronted with recent advances in understanding the full complexity of 3D genome organization, its cell-type dependence and its influence on gene regulation [61, 62].

Our investigation demonstrates a link between genetic risk and 3D genome partitioning into topologically-insulated domains. A genetic variation located in TAD borders may weaken the insulation of adjacent TADs, which prevents spurious interactions between genes and enhancers located in adjacent TADs. The larger frequency of cancers among these diseases support the importance of TAD border weakening in a subset of cancers, and emphasizes the different type of gene mis-regulation involved in cancer etiology. A cancer generally involves the malfunction of numerous genes, which is more readily achieved by the extended deregulations induced by weakening of a single TAD border rather than affecting each gene individually. TAD disruption induced by somatic mutations has been observed in cancers [31], and our results suggest that an at-risk variant SNPs may act as a head start.

Our results offer the proof-of-concept of a novel criterion for filtering SNPs according to their 3D genomic location, and identifying especially relevant associations, i.e. SNP prioritization. Our study opens a new research avenue in the personalized diagnosis of genetic risk, based on the interplay between 3D genome organization and the location of at-risk SNPs. Dissecting the functional correlates of the preferential location of risk loci in TAD borders now challenges experimental studies. Various experiments, including genome editing to monitor the allelic form of specific loci and chromosome conformation capture techniques, could bring a mechanistic support to this novel statistical evidence of a link between 3D genome organization and the risk of developing certain complex diseases.

## Acknowledgments

The authors thank Nastasija Mijovic, Cosette Rebouissou, Marina Villaverde, Valentin Ruault and the members of the CNRS GDR 3536 ‘ADN’ for insightful discussions.

## Author contributions

AL and MTH designed the study.

KPJ and LC performed the bioinformatic analysis.

KPJ performed the statistical analysis and devised the integrated pipeline.

KPJ and MTH performed the network analysis and prepared the figures.

All authors contributed to the interpretation of the results.

All authors edited the manuscript and approve the final version.

## Abbreviations

3D: 3-dimensional
GWAS: genome-wide association studies
hESC: human embryonic stem cell
Hi-C: genome-wide chromosome conformation capture
IMR90: human fetal lung fibroblasts
SNP: single-nucleotide polymorphism
daSNP: disease-associated SNP
TAD: topologically associating domain

## SUPPORTING INFORMATION

**S1 Fig.**
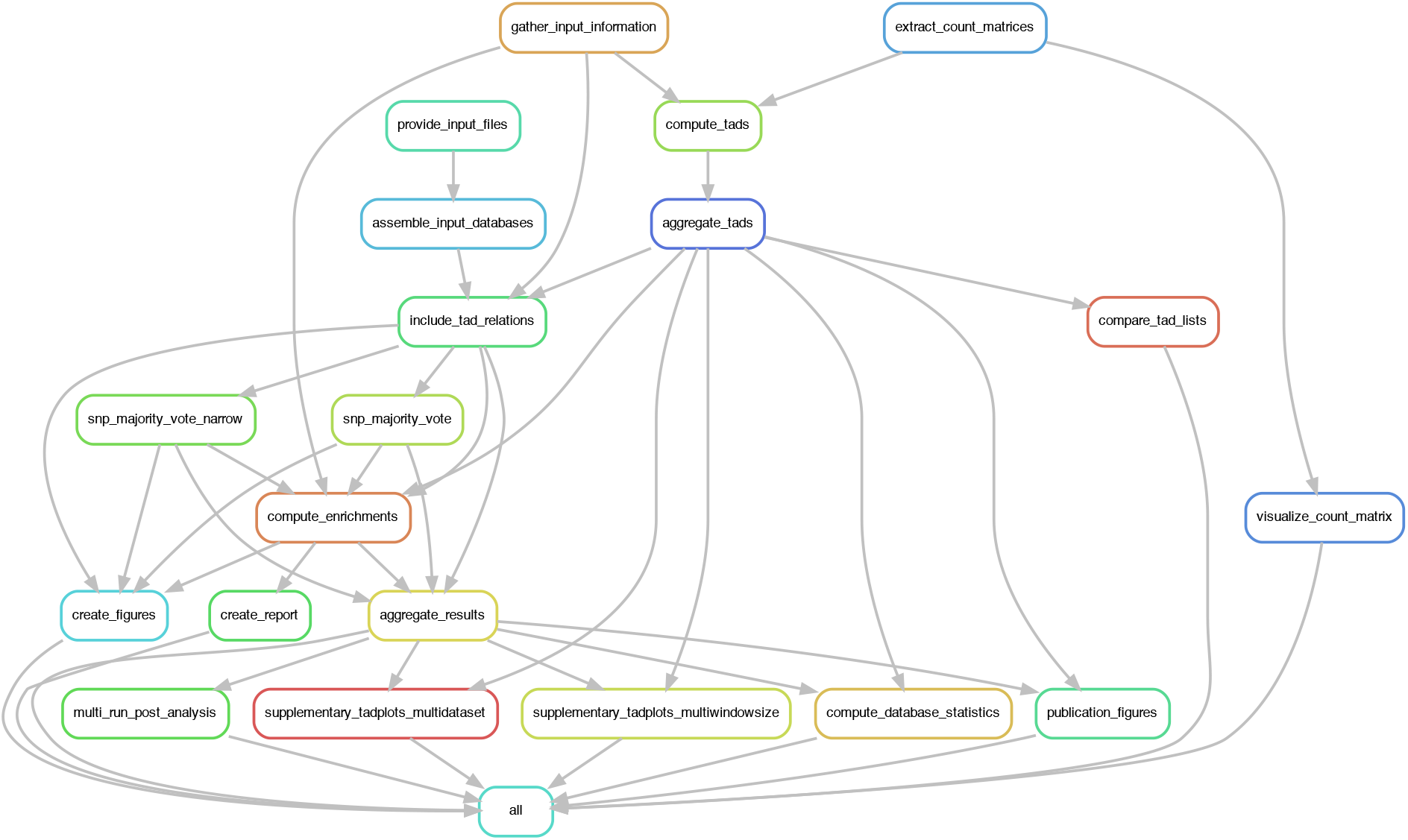
Rule graph of the integrated pipeline. The figure displays an automated drawing of the bioinformatic and statistical pipeline devised for the study. For each disease, the potential TAD border enrichment in disease-associated SNPs is assessed from the associations available in the GWAS catalog and Hi-C data, downloaded from ftp://cooler.csail.mit.edu/coolers. The test involves the determination of TADs using TopDom algorithm [S1] and the computation of the *p-*value quantifying the statistical significance of the enrichment (see *Materials and methods)*. The subsequent steps of the analysis, for instance the aggregation of the results over several values of the parameter *k* of the TAD caller, the plots of enrichment histograms (Fig 2) or percentages (Fig 3), have been integrated in the pipeline, available at: https://github.com/kpj/GeneticRiskAndTADs. The whole analysis can thus be implemented straightforwardly for any Hi-C dataset in .cool format.

**S2 Fig.**
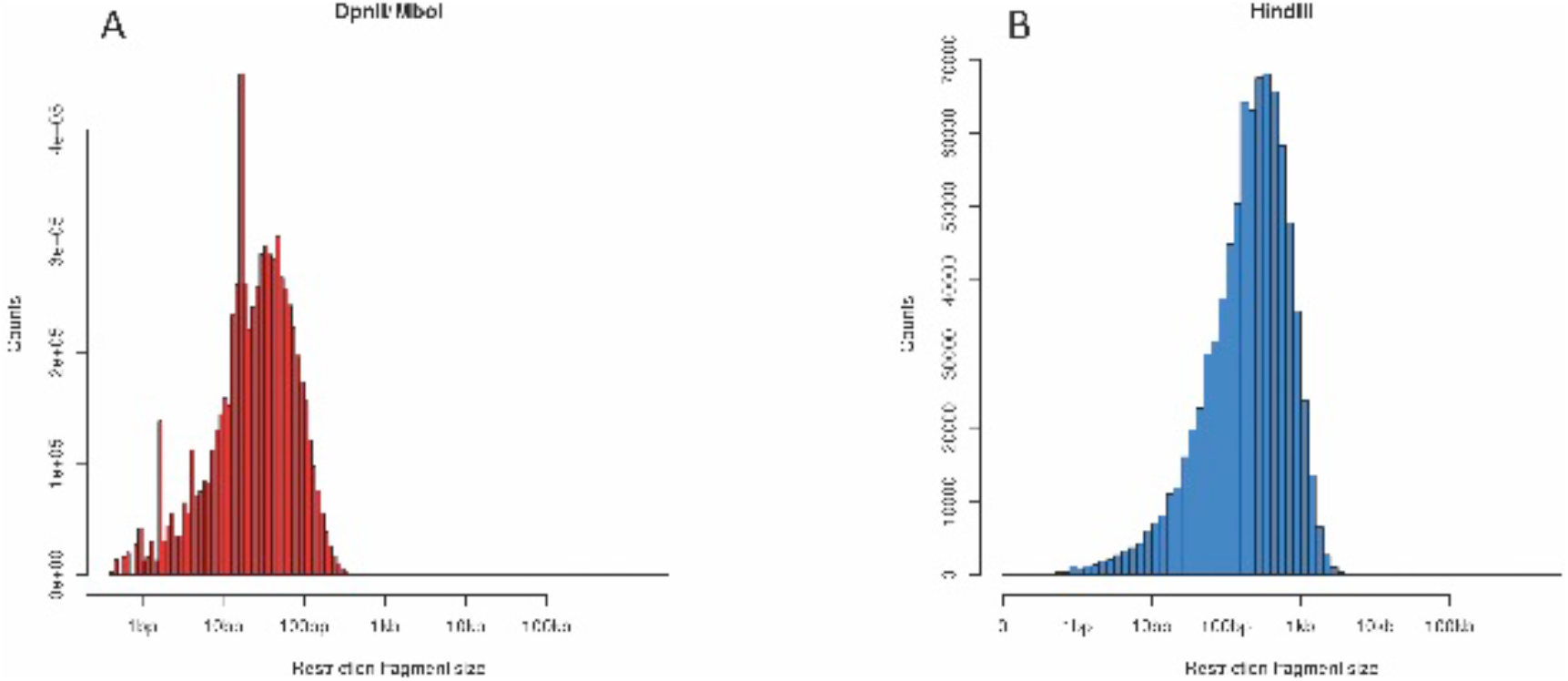
Restriction fragment size distributions. The distribution of the fragment size (size in log scale on the horizontal axis) has been determined from the positions of the restriction sites on the genome (using *digest_genome* script from HiC-Pro toolbox [S2]) on the NCBI genome version) for: (A) the restriction enzymes MboI or DpnII, producing the same fragments (the enzymes recognize the same sequence) and (B) the restriction enzyme HindIII producing larger fragments (note the different scale for the counts on the vertical axis). These distributions show that all restriction fragments obtained with MboI or DpnII have a size below 1kb, whereas a non-negligible percentage of DNA fragments obtained with HindIII have a size larger than 1kb, however below 10kb, indicating that Hi-C resolution is mostly limited by sequencing depth.

**S3 Fig.**
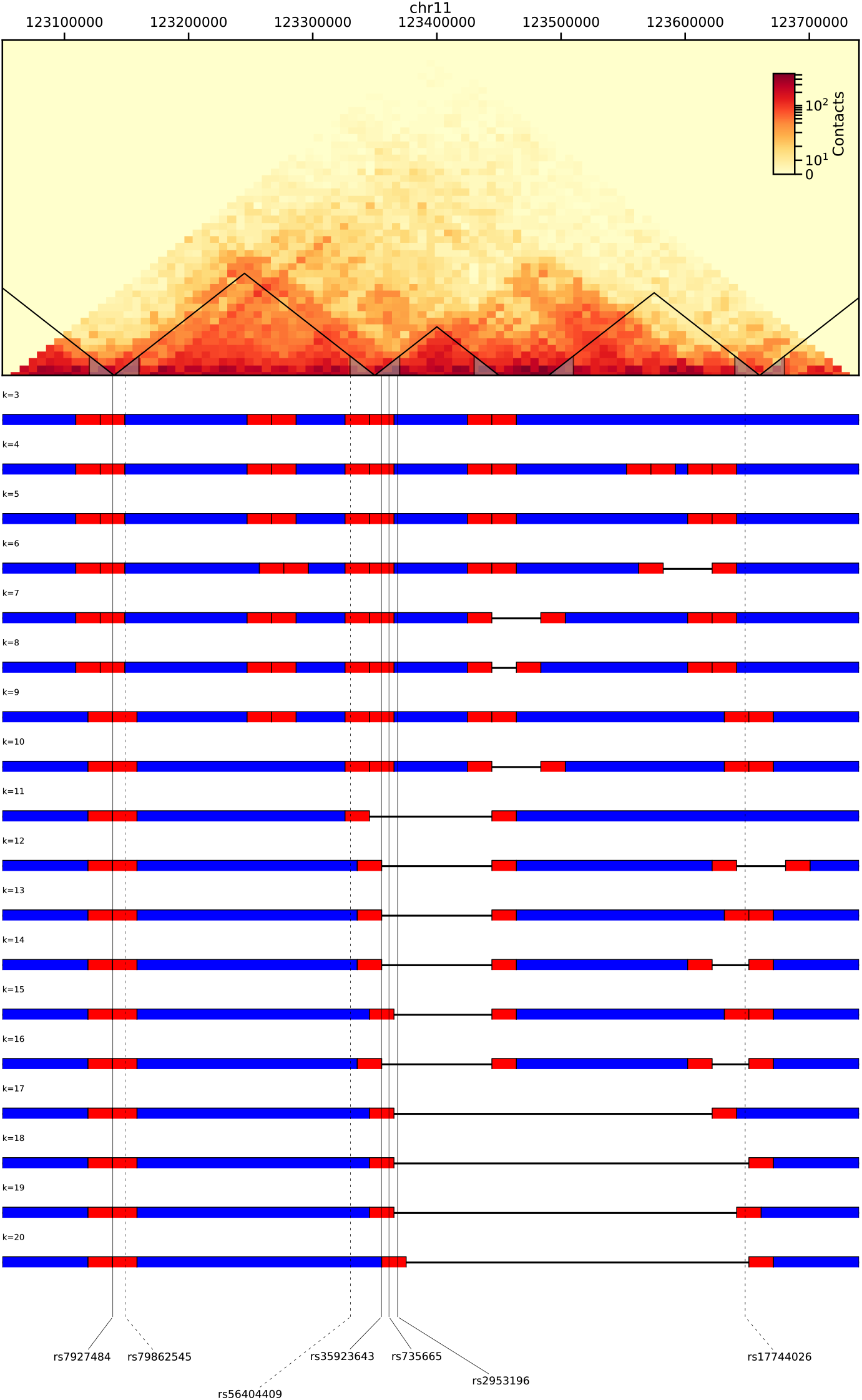
Variation of TADs and TAD borders at varying TopDom parameter *k*. The top panel displays the Hi-C contact matrix as a heat map (see the color bar, the redder the more contacts), here for a region of chr11 (chr11: 123050000-123750000, hg19 coordinates, embedding the region in Fig 1), drawn from Hi-C data published in [S3], for IMR90 cell type, at 10kb-resolution. TADs determined with TopDom, for a window size *k*=10, are underlined together with their internal 20kb-borders. Vertical lines pinpoint SNPs located in a TAD border (full line for cancer-associated SNPs, dashed lines for SNPs associated with non-cancer diseases). TADs obtained at increasing value of *k*, from *k*=3 to *k*=20, are schematically displayed in the lines below (blue: TAD body, red: TAD border, value of k indicated above each line at its left end).

**S4 Fig.**
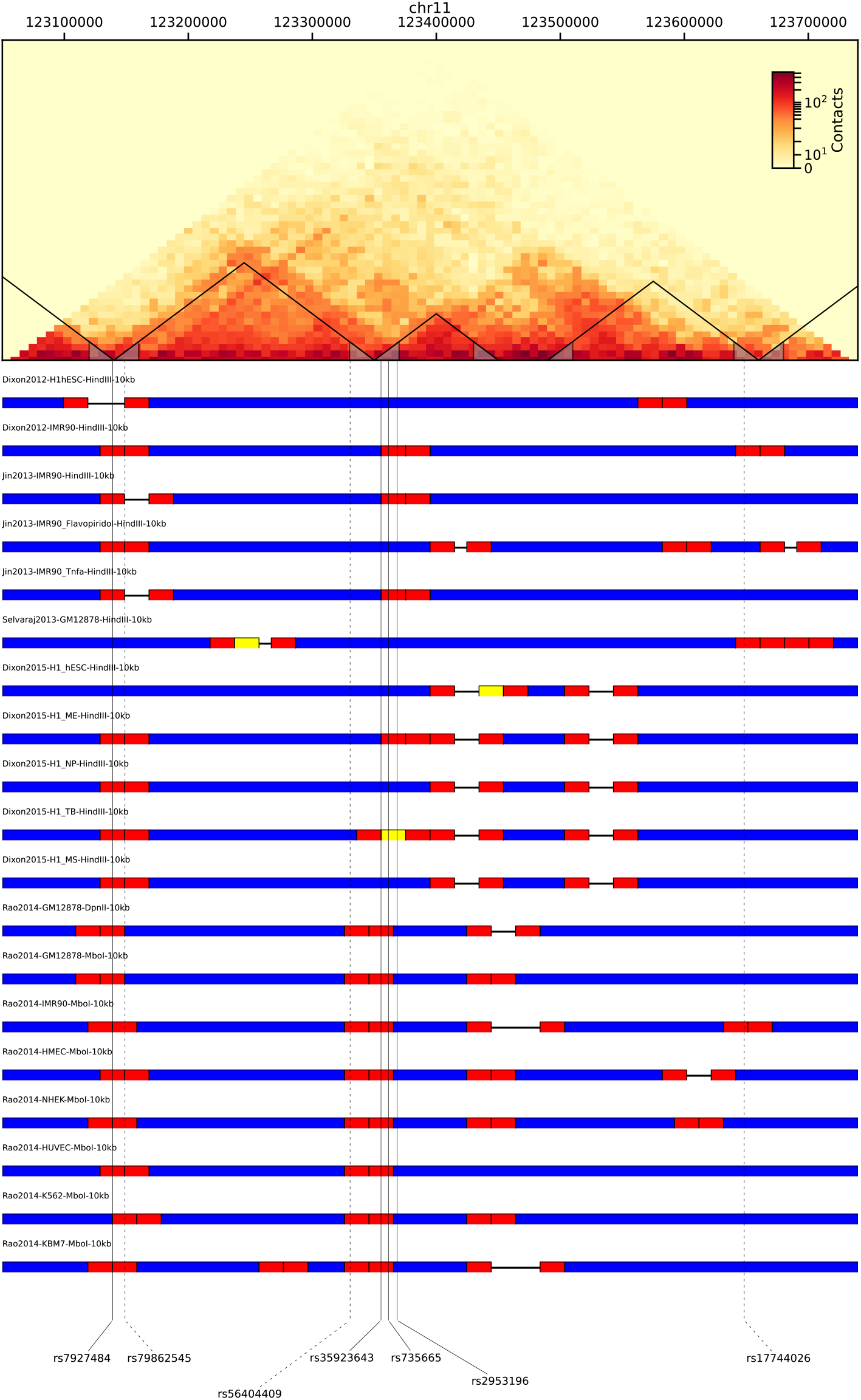
Variation of TAD and TAD borders across data sources. Same as the previous S3 Fig, now for different data sources, indicated above each line, at a fixed value *k*=10 of TopDom window parameter (blue: TAD body, red: TAD border defined as the end region of size 20kb within the TAD, yellow: situations where a TAD is too short and its two borders overlap). The Hi-C map is based on Hi-C data from [S3], IMR90 cell type, at 10kb-resolution, for the region chr11: 123050000-123750000, hg19 coordinates (same region as in S3 Fig and embedding the region in Fig 1).

**S5 Fig.**
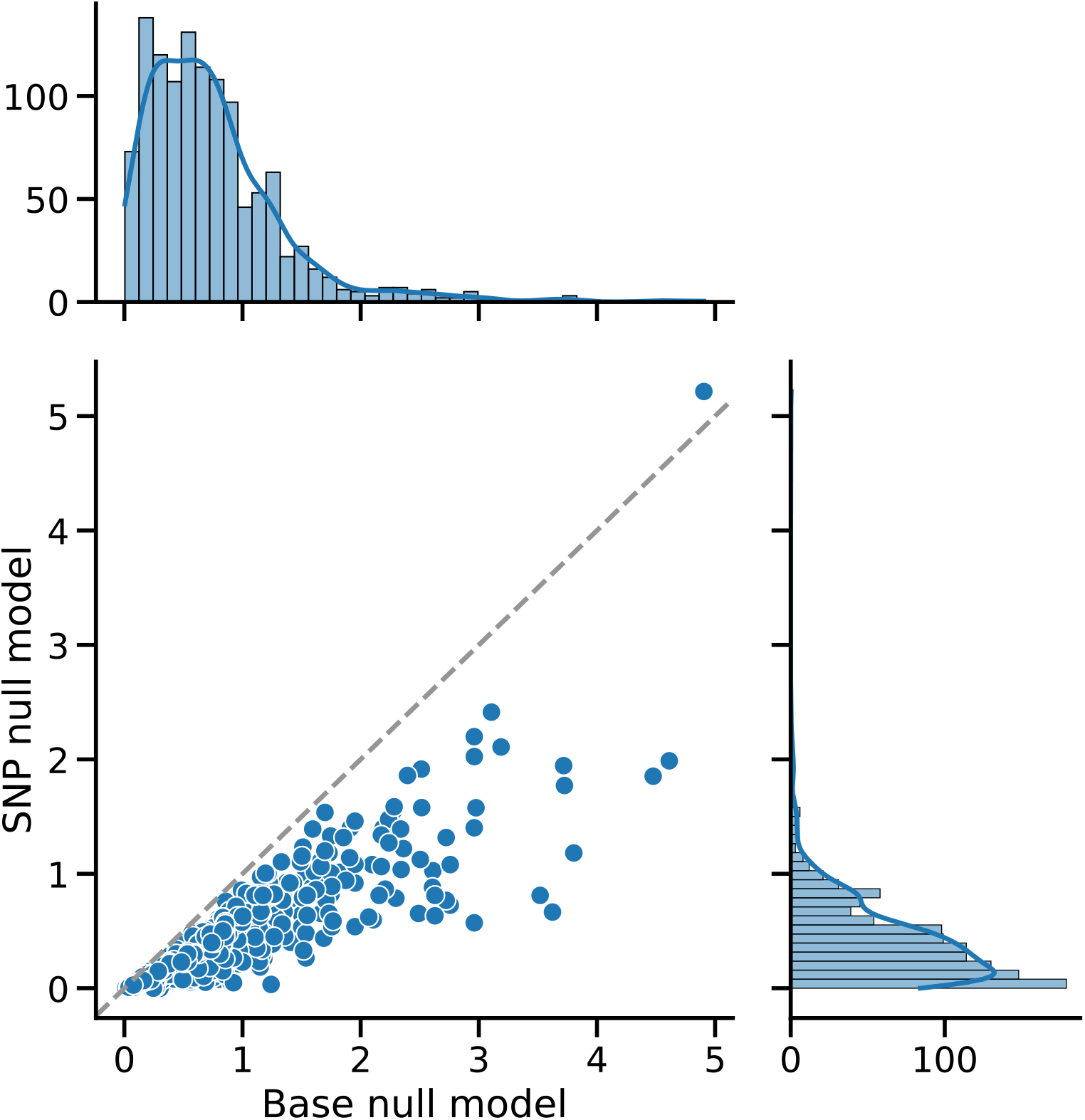
Comparison of two null models for assessing TAD border enrichment in daSNPs. The figure presents a scatter plot of the enrichment statistical significance [-log(*p-*value)], one dot per disease, for two different null models (see *Materials and methods*) and, on the right and top respectively, the *p-*value distribution over all diseases for each null model. The first null model (horizontal axis in the scatter plot, distribution on top) is defined for each disease as a homogeneous distribution of the associated SNPs along the genome, at the base pair level, whereas the second one (vertical axis in the scatter plot, distribution on the right) is defined for each disease as a homogeneous distribution of the associated SNPs within the ensemble of all disease-associated SNPs gathered from the GWAS catalog (not including SNPs associated with non-pathological traits). The *p-*value assessing for each disease the over-representation of its associated SNPs in TAD borders is computed using the cumulative hypergeometric distribution, then corrected for multiple testing using the Benjamini-Hochberg procedure [S4] separately for cancers and non-cancer diseases (see *Materials and methods)*. The correlation coefficient between the results obtained with each null model is 0.92. Hi-C data from [S3], IMR90 cell type, 10kb resolution; TAD determination with TopDom window parameter value *k*=10.

**S6 Fig.**
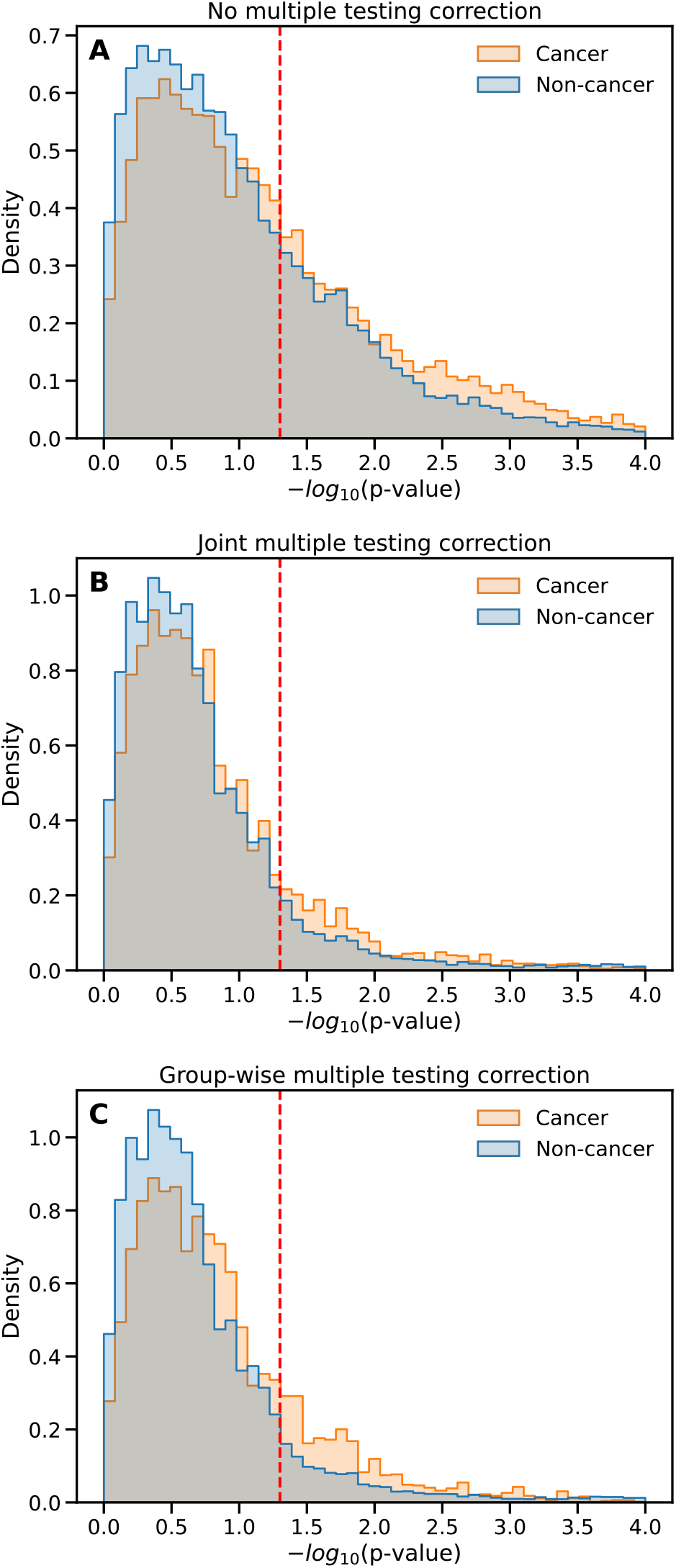
Multiple testing correction in assessing TAD border enrichment in daSNPs. Histograms of [-log(*p-*value)] for cancers (orange) and non-cancer diseases (blue, overlap in grey) (A) without multiple-testing correction; (B) with a correction applied to all diseases jointly; (C) with a correction applied separately to cancers and non-cancer diseases (group-wise correction, see Fig 2). In cases B and C, the correction followed the Benjamini-Hochberg procedure [S4] controlling the false discovery rate. The significance threshold at 5% is indicated by the dashed red line. Histograms are normalized separately for cancers and non-cancer diseases. Same underlying Hi-C data and setting as in Fig 2.

**S7 Fig.**
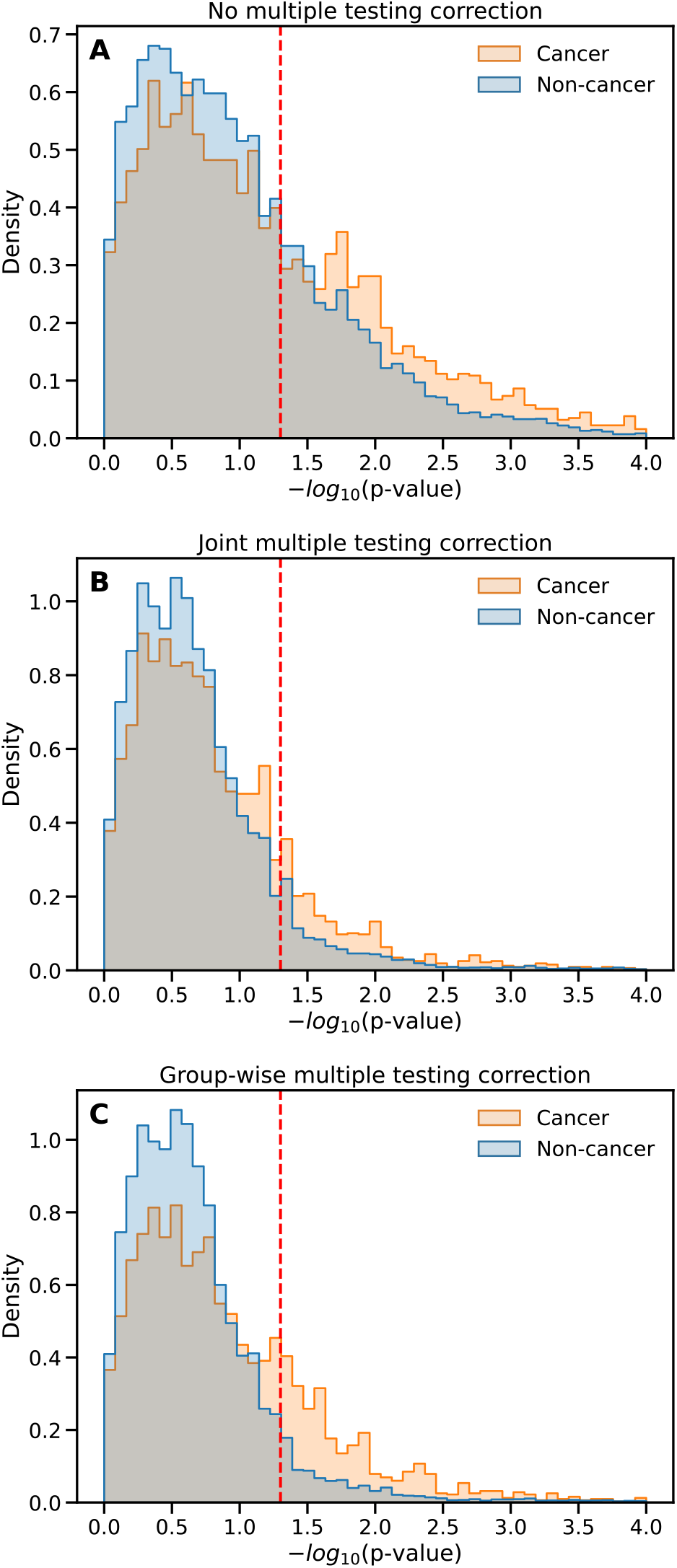
Multiple testing correction in assessing TAD border enrichment in intergenic daSNPs. Same as S6 Fig considering intergenic daSNPs only. The figure displays the normalized histograms of [-log(*p-* value)] for cancers (orange) and non-cancer diseases (blue, overlap in grey) and (A) without multiple-testing correction; (B) with a correction applied to all diseases jointly; (C) with a correction applied separately to cancers and non-cancer diseases (group-wise correction, see Fig 2).

**S8 Fig.**
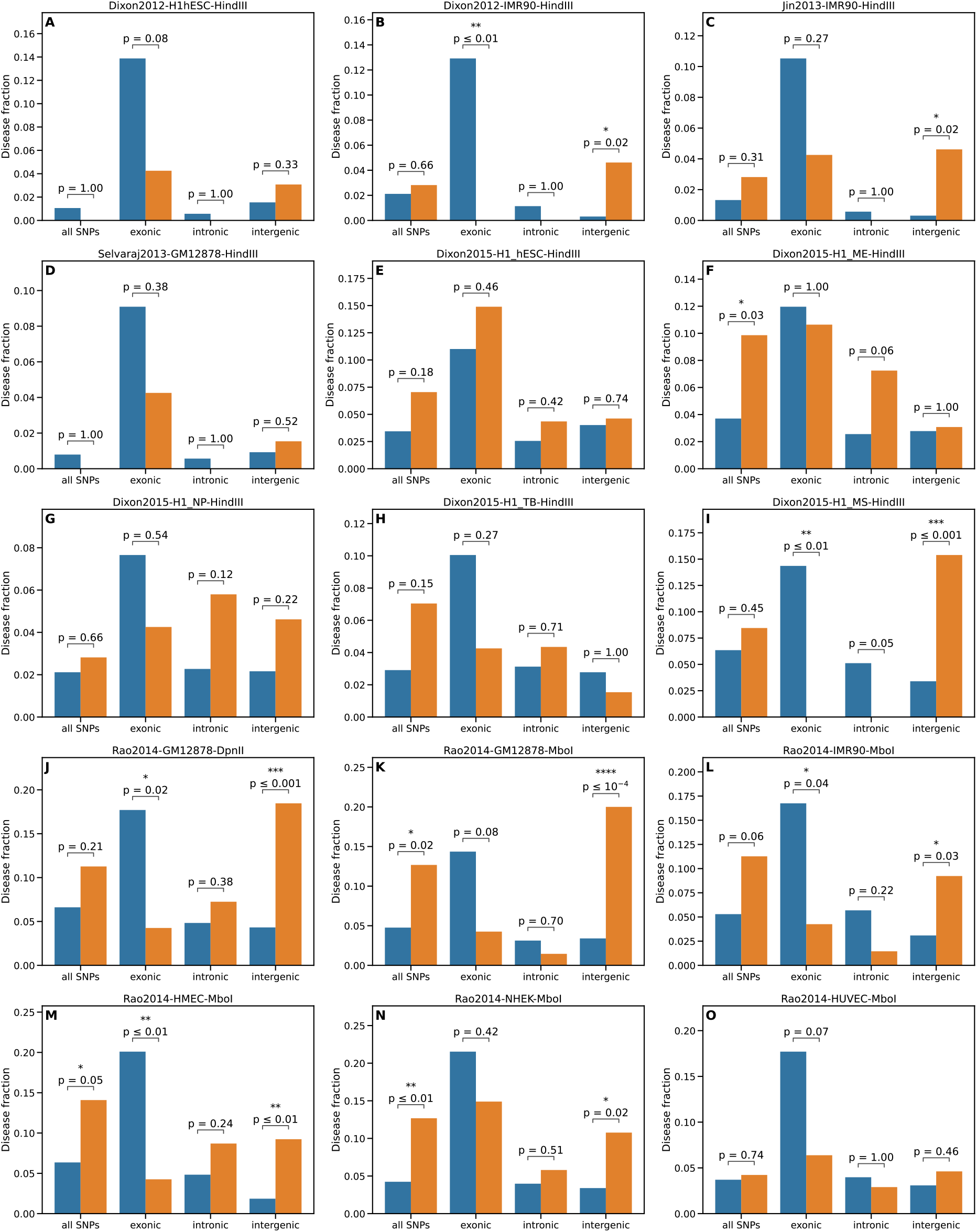
TAD border enrichment across data sources. The relative dominance of cancers among the diseases displaying a preferential location of their associated SNPs in TAD borders is investigated across Hi-C datasets (same setting as in Fig 3). The datasets were obtained in different laboratories (main ordering of the panels), or/and with different restriction enzymes (HindIII, MboI or DpnII, as indicated in the caption, see S2 Fig) or/and in different cell types: human embryonic stem cells (H1 hESC) and derived cell lines: mesendoderm (H1_ME), neural progenitors (H1_NP), trophoblast-like cells (H1_TB) and mesenchymal cells (H1_MS); human lymphoblastoid cell line (GM12878); fetal lung fibroblasts of Caucasian origin (IMR90); human mammary epithelial cells (HMEC); normal human epidermal keratinocytes (NHEK). human umbilical vein endothelial cells (HUVEC) [S3, S5-S8].

**S1 File: Typical numbers of diseases and SNPs involved in the analysis**

About 350 Mb are located within TAD borders, representing about 12% of the genome and 13% of the number of base pairs located in TADs. However, this number of base pairs located in TAD borders varies with the value of the parameter *k* used in the TAD caller TopDom, roughly decreasing when *k* increases. Exact values in the case of IMR90 cell type, data from [S3], are given in S1 Table.

**S1 Table:** Genome fraction located in TAD borders. The fraction of the genome (resp. of the total number of base pairs in TADs) located in TAD borders is given for different values of the parameter *k* (window size) of the TAD caller TopDom (underlying Hi-C data from [S3 27], IMR90 cell type).

| Window size $k$ | TAD border contents relative to genome | TAD border contents relative to TADs |
| --- | --- | --- |
| 3 | 12.8% | 14.4% |
| 4 | 14.1% | 15.8% |
| 5 | 14.3% | 16% |
| 6 | 14.1% | 15.7% |
| 7 | 13.6% | 15.2% |
| 8 | 12.9% | 14.4% |
| 9 | 12.3% | 13.7% |
| 10 | 11.7% | 13.1% |
| 11 | 11% | 12.3% |
| 12 | 10.5% | 11.8% |
| 13 | 10.1% | 11.3% |
| 14 | 9.7% | 10.8% |
| 15 | 9.2% | 10.3% |
| 16 | 8.9% | 10% |
| 17 | 8.6% | 9.6% |
| 18 | 8.3% | 9.3% |
| 19 | 8.1% | 9% |
| 20 | 7.8% | 8.7% |

449 EFOs have been considered in the study, among which 71 cancers (that is, 16%)

The number of disease-associated SNPs considered in the study (without multiplicity) is 21,183 among which about 2800 (13%) are located in TAD borders. The overall number of base pairs located in TAD borders and border SNP counts also vary with the data source and the resolution (10kb or 50kb) of the contact map, as described in S2 Table.

The number of cancer-associated SNPs is 3,319 among which about 470 (14%) are located in TAD borders

8,438 (40%) disease-associated SNPs are intergenic

1,058 are intergenic and located in TAD borders

1,275 are intergenic and associated with cancer

176 are intergenic, associated with cancer and located in TAD borders

11,552 (55%) disease-associated SNPs are intronic

1,508 are intronic and located in TAD borders

1,854 are intronic and associated with cancer

257 are intronic, associated with cancer and located in TAD borders

1,171 (5%) disease-associated SNPs are exonic

223 are exonic and located in TAD borders

185 are exonic and associated with cancer

35 are exonic, associated with cancer and located in TAD borders

**S2 Table:** Genome fraction and number of SNPs located in TAD borders for different datasets. The fraction of the genome (resp. of the total number of base pairs in TADs) and the number (resp. the fraction) of disease-associated SNPs located in TAD borders is given for different data sources. Unless otherwise stated, data from [S3] were obtained from experiments using MboI enzyme, whereas all other datasets were obtained from experiments using HindIII enzyme.

| Data set | TAD border contents<br>w.r.t. genome | TAD border contents<br>w.r.t. all TADs | border SNP<br>count | fraction of border SNPs<br>w.r.t. number of daSNPs |
| --- | --- | --- | --- | --- |
| Dixon et al 2012<br>H1 hESC<br>HindIII enzyme | 8.7% | 9.7% | 1969 | 9.3% |
| Dixon et al 2012<br>IMR90<br>HindIII enzyme | 8.8% | 9.9% | 2018 | 9.5% |
| Jin et al 2013<br>IMR90<br>HindIII enzyme | 7.7% | 8.6% | 1834 | 8.7% |
| Selvaraj 2013<br>GM12878<br>HindIII enzyme | 12.2% | 13.7% | 2552 | 12% |
| Dixon et al 2015<br>H1 hESC,<br>HindIII enzyme | 13.1% | 14.7% | 3179 | 15% |
| Dixon et al 2015<br>H1_ME,<br>HindIII enzyme | 13.6% | 15.3% | 3223 | 15% |
| Dixon et al 2015<br>H1_NP<br>HindIII enzyme | 14.4% | 16.1% | 3398 | 16% |
| Dixon et al 2015<br>H1_TB<br>HindIII enzyme | 12.8 | 14.3% | 3087 | 15% |
| Dixon et al 2015<br>H1_MS<br>HindIII enzyme | 12.6% | 14% | 3053 | 14% |
| Rao et al 2014<br>GM12878<br>DpnII enzyme | 11.7% | 13.1% | 3056 | 14% |
| Rao et al 2014<br>GM12878<br>MboI enzyme | 10.8% | 12% | 2789 | 13% |
| Rao et al 2014<br>IMR90<br>MboI enzyme | 11.7% | 13.1% | 3034 | 14% |
| Rao et al 2014<br>HMEC<br>MboI enzyme | 13.1% | 14.7% | 3408 | 16% |
| Rao et al 2014<br>NHEK<br>MboI enzyme | 15% | 16.8% | 3698 | 17% |
| Rao et al 2014<br>HUVEC<br>MboI enzyme | 11.8% | 13.2% | 3005 | 14% |

**S3 Table:** Values of network coherence for various subgraphs in SNP-based diseasome networks. Nodes in the diseasomes represent diseases, distinguishing cancers or non-cancer diseases. Four SNP-based diseasome networks have been constructed, depending on the meaning of an edge between two nodes. In the network labelled ‘border’, an edge is drawn between two diseases when they share at least one border SNP, i.e. a SNP located in a TAD border for a majority of values of TopDom parameter *k*. Non-border SNPs are defined as the complementary set of SNPs, yielding the network labelled ‘non-border’. The networks labelled ‘border intergenic’, and ‘non-border intergenic’ are obtained when the additional condition of being intergenic is imposed on the shared SNPs. Note that these graphs are not disjoint; two diseases can be linked in two or more of these diseasome networks. The level of clustering (density of internal links) of various subgraphs, compared to a situation of random wiring, has been estimated using the notion of network coherence, defined as the fraction of connected nodes within the subgraph, z-transformed with a null model of randomly drawn node sets of the same size (see *Materials and methods*). The considered subgraphs correspond to sets of diseases: cancers, non-cancer diseases, cancers whose associated SNPs are preferentially located in TAD borders for a majority of values of *k* (‘enriched cancer’) or not (‘not enriched cancer’) and similar definitions for enriched and not enriched non-cancer diseases. Network coherence provides an absolute quantification, here computed for the 6 subgraphs in the four diseasome networks. A random subgraph would have a vanishing network coherence, a positive value indicates a high internal connectivity while a negative value reveals that the subgraph is less connected than a random set of nodes in the overall network.

| diseasome network | cancer | non-cancer | enriched cancer | enriched non-cancer | not enriched cancer | not enriched non-cancer |
| --- | --- | --- | --- | --- | --- | --- |
| ‘border’ | **4.02** | **-1.26** | **2.94** | **2.09** | **3.20** | **-1.50** |
| ‘border intergenic’ | **2.75** | **-0.15** | **4.24** | **4.04** | **1.67** | **-0.60** |
| ‘non-border’ | **5.02** | **-1.59** | **3.45** | **3.55** | **4.56** | **-2.92** |
| ‘non-border intergenic’ | **2.62** | **0.09** | **2.29** | **4.85** | **2.61** | **-1.29** |

